# Determining The Structure of the Bacterial Voltage-gated Sodium Channel NaChBac Embedded in Liposomes by Cryo Electron Tomography and Subtomogram Averaging

**DOI:** 10.1101/2023.04.24.538027

**Authors:** Shih-Ying Scott Chang, Patricia M. Dijkman, Simon A. Wiessing, Misha Kudryashev

## Abstract

Voltage-gated sodium channels shape action potentials that propagate signals along cells. When the membrane potential reaches a certain threshold, the channels open and allow sodium ions to flow through the membrane depolarizing it, followed by the deactivation of the channels. Opening and closing of the channels is important for cellular signalling and regulates various physiological processes in muscles, heart and brain. Mechanistic insights into the voltage-gated channels are difficult to achieve as the proteins are typically extracted from membranes for structural analysis which results in the loss of the transmembrane potential. Here, we report the structural analysis of a bacterial voltage-gated sodium channel, NaChBac, reconstituted in liposomes under an electrochemical gradient by cryo electron tomography and subtomogram averaging. We show that the small channel, most of the residues of which are embedded in a membrane, can be localized using a genetically fused GFP. GFP can aid the initial alignment to an average resulting in a correct structure, but does not help for the final refinement. At a moderate resolution of ∼16 Å the structure of NaChBac in an unrestricted membrane bilayer is 10% wider than the structure of a purified protein previously solved in nanodiscs, suggesting the potential movement of the peripheral voltage-sensing domains. Our structural analysis explores the limits of structural analysis of membrane proteins in membranes.

**HIGHLIGHTS:**

- Structural analysis of the bacterial voltage-gated sodium channel NaChBac in lipid vesicles under the resting membrane potential by cryo electron tomography and subtomogram averaging.
- Fused GFP allows identification of a 120-kDa mostly transmembrane protein in tomograms, and helps for the initial alignment but not for the final refinements.
- The map of NaChBac in liposomes at a resolution of 16.3 Å is ∼10% wider than the protein structure in a nanodisc.

## INTRODUCTION

Biological membranes provide physical barriers in cells and organelles, and allow the maintenance of electrochemical gradients that can be used to trigger various membrane proteins for life-critical functions (Watson, 2015; Sundelacruz et al, 2009). Voltage-gated ion channels (VGICs) are a class of transmembrane proteins that are selectively permeable to ions such as Na^+^, K^+^, Ca^2+^, and Cl^-^ when they are activated and undergo conformational changes due to changes in membrane potential around the channels (Bezanilla, 2008). In neuronal signaling and muscular contraction, Na^+^ influx through voltage-gated sodium channels (VGSCs) corresponds to the rapid rising phase of the action potential in the membranes of neurons and other electrically excitable cells (Hodgkin & Huxley, 1939, 1945), and VGSCs show a series of conformational changes between open, closed, and inactivated states during a cycle of an action potential (Yarov-Yarovoy et al, 2012; Lenaeus et al, 2017). Each action potential is followed by a refractory period, during which the VGSCs enter an inactivated state while the Na^+^ and K^+^ ions return to their resting state distributions across the membrane. In most neurons the resting potential is approximately -70 mV, and VGSCs have transitioned back to their resting state and have re-established the process for the next action potential.

NaChBac is a model bacterial VGSC from *Bacillus halodurans* widely used to study their structure and function. Each monomer consists of 6-transmembrane helices, helices S1-S4 form voltage-sensing domains for each unit, while S5 and S6 form the common pore domain with the other units (Charalambous & Wallace, 2011) to have an overall tetrameric scaffold. X-ray crystallography and recent cryo electron microscopy (cryo-EM) single particle analysis (SPA) provided mechanistic insights into the activation sequence of VGICs. Crystal structures of the prokaryotic Na_v_Ms from *Magnetococcus marinus* were determined in an open state, with an open selectivity filter leading to an open activation gate at the intracellular membrane surface to allow hydrated sodium ions to pass through (McCusker et al, 2012; Sula et al, 2017). The crystal structure of Na_v_Ab from *Arcobacter butzleri* was captured in a closed-pore conformation by amino acid substitutions (Payandeh et al, 2011), and the cryo-EM structure of NaChBac in nanodiscs showed a potentially inactivated state (Gao et al, 2020). For eukaryotic VGSCs, cryo-EM structures were obtained of Na_v_1.4 in complex with the β1 subunit from electric eel and human in an open pore state (Yan et al, 2017; Pan et al, 2018), and of Na_v_PaS from American cockroach in multiple closed conformations (Shen et al, 2017). In order to stabilize the resting conformation of VGSCs for cryo-EM structure determination, disulfide crosslinking in the voltage-sensing module was used for Na_v_Ab (Wisedchaisri et al, 2019). Also, tarantula toxin and voltage-shifting mutations were designed for trapping the resting state of Na_v_1.7 chimera (Wisedchaisri et al, 2021). A structure of the Eag potassium channel under membrane potential has been recently reported, describing conformational changes in the voltage sensing domain and the interplay between the channel and lipids (Mandala & MacKinnon, 2022). For smaller VGSCs it is still a challenge to have a system which can allow structural analysis in unrestricted membranes and in presence of the electrochemical gradients. A structure of VGSCs under a physiological resting membrane potential to provide support for the classical ‘‘sliding helix’’ model for gating from the resting state to the activated state is still missing (Wisedchaisri et al, 2019).

Proteoliposomes, which are lipid vesicles with reconstituted membrane proteins, provide an excellent system for functional and structural studies of membrane proteins under conditions that mimic those *in vivo* (Sejwal et al, 2017). Proteoliposomes allow preservation of the functional lipid environment, generation of transmembrane ionic gradients and do not restrict the motion of transmembrane helices in the membrane plane. Despite extensive functional characterizations using proteoliposomes, successful attempts to employ this system for structural elucidation of membrane proteins are still limited. Structures of hBK channel (Wang & Sigworth, 2009; Tonggu & Wang, 2022), AcrB transporter (Yao et al, 2020), Eag K_v_ channel (Mandala & MacKinnon, 2022), and PIEZO1 channel (Yang et al, 2022) in liposomes have been determined by single particle cryo-EM (SPA) at resolutions ranging from 3.5 to 7 angstrom. However, without a significantly large soluble domain, it may be challenging to “identify” fully transmembrane proteins in electron micrographs for alignment and averaging (Yao et al, 2020). Indeed, the molecular weight of these proteins for which structures were solved from proteoliposomes is relatively high: hBK channel is ∼500 kDa, AcrB transporter is ∼350 kDa, Eag K_v_ channel is ∼390 kDa, and Piezo1 channel is ∼860 kDa, and they have substantial extramembranous domains.

Cryo electron tomography (cryo-ET) and subtomogram averaging (StA) has also been used for determining structures of membrane proteins in lipid vesicles: structures of the small membrane protein MspA (∼160 kDa) and the serotonin receptor ion channel 5-HT_3_R (∼275 kDa) were determined at resolutions of ∼17 and ∼12 A respectively in 2012 and 2015 (Eibauer et al, 2012; Kudryashev et al, 2016). Later, a structure of a large ion channel RyR1 was reported *in situ* first at 12.6 Å resolution and then at subnanometer resolution allowing resolving alpha helices (Chen & Kudryashev, 2020; Sanchez et al, 2020). While the resolution of StA structures is lower than typically obtained by the single particle reconstructions, we hypothesized that StA might be able to target smaller membrane proteins in lipid vesicles, as tomograms contain the third dimension compared to 2D imaging in SPA. Here, we attempted to capture the resting state of the small (∼120 kDa) bacterial VGSC NaChBac by cryo electron tomography. To this end, we purified NaChBac, reconstituted it into liposomes, introduced a transmembrane potential to this system, and performed structural analysis by cryo-ET and subtomogram averaging.

## RESULTS

### Design, sample preparation and reconstitution into proteoliposomes

We produced His_6_-GFP^A206K^-NaChBac in *Escherichia coli* C41(DE3) and purified it using a two-step purification approach of affinity and size-exclusion chromatography (SI Appendix, Fig. S1 A to C). GFP was fused to the N-terminal end of each of the channel subunits to function as an “anchor” for identifying the channels in the membrane. We next reconstituted NaChBac into proteoliposomes using *E. coli* polar lipid extract to mimic the native bacterial membrane environment. A relatively high lipid-to-protein ratio of 2:1 (weight to weight) was used in order to maximise the number of particles per vesicle and thus the efficiency of data collection. For the reconstitution, the detergent DDM was removed by dialysis and followed by Bio-beads (Rigaud et al, 1998). We found that reducing the rate of detergent removal improved the reconstitution efficiency, resulting in more protein copies per proteoliposome as observed by cryo-EM. However, due to the very low critical micelle concentration of DDM, a relatively long time for the dialysis (around one week) and subsequent incubation with Bio-beads was necessary, otherwise the resulting proteoliposomes showed the membrane with a “fluid edge” (SI Appendix, Fig. S2 B). Gradient ultracentrifugation showed a significant protein band shift for the proteoliposomes compared to empty liposomes, and the 55-kDa his-tagged GFP^A206K^-NaChBac showed no signs of contamination or protein degradation by SDS-PAGE (Fig. 2 A).

In order to generate a membrane potential we used the well-established protocol of resuspending proteoliposomes prepared with a buffer with a high KCl concentration into a buffer with a low KCl concentration followed by addition of a potassium ionophore valinomycin. The membrane potential across the NaChBac-containing proteoliposome bilayer was assayed using the voltage-sensitive cationic fluorescent dye JC-1 (Reers et al, 1991; Chanda & Mathew, 1999) (Fig. 1 A). Proteoliposomes were prepared in a buffer with 150 mM KCl and were resuspended into 3 mM KCl buffer in the presence of 1 μM valinomycin (DMSO as a control experiment), (Fig. 2 B) resulting in a membrane potential of -100 mV at room temperature (298 K) as determined using the Nernst equation for the potassium diffusion potential calculation (Smiley et al, 1991) (Fig. 1 D). Measuring the fluorescence over time showed stable levels of membrane potential over tens to thousands of seconds. Increasing the external concentration of KCl led to a decrease in JC-1 fluorescence due to less membrane negative potential, indicating the K^+^-selective permeability and the stability of proteoliposomes (Fig. 1 B and C). These polarized vesicles at the negative resting membrane potential (-100 mV) condition were immediately applied to a holey gold grid and frozen.

**Figure 1.**
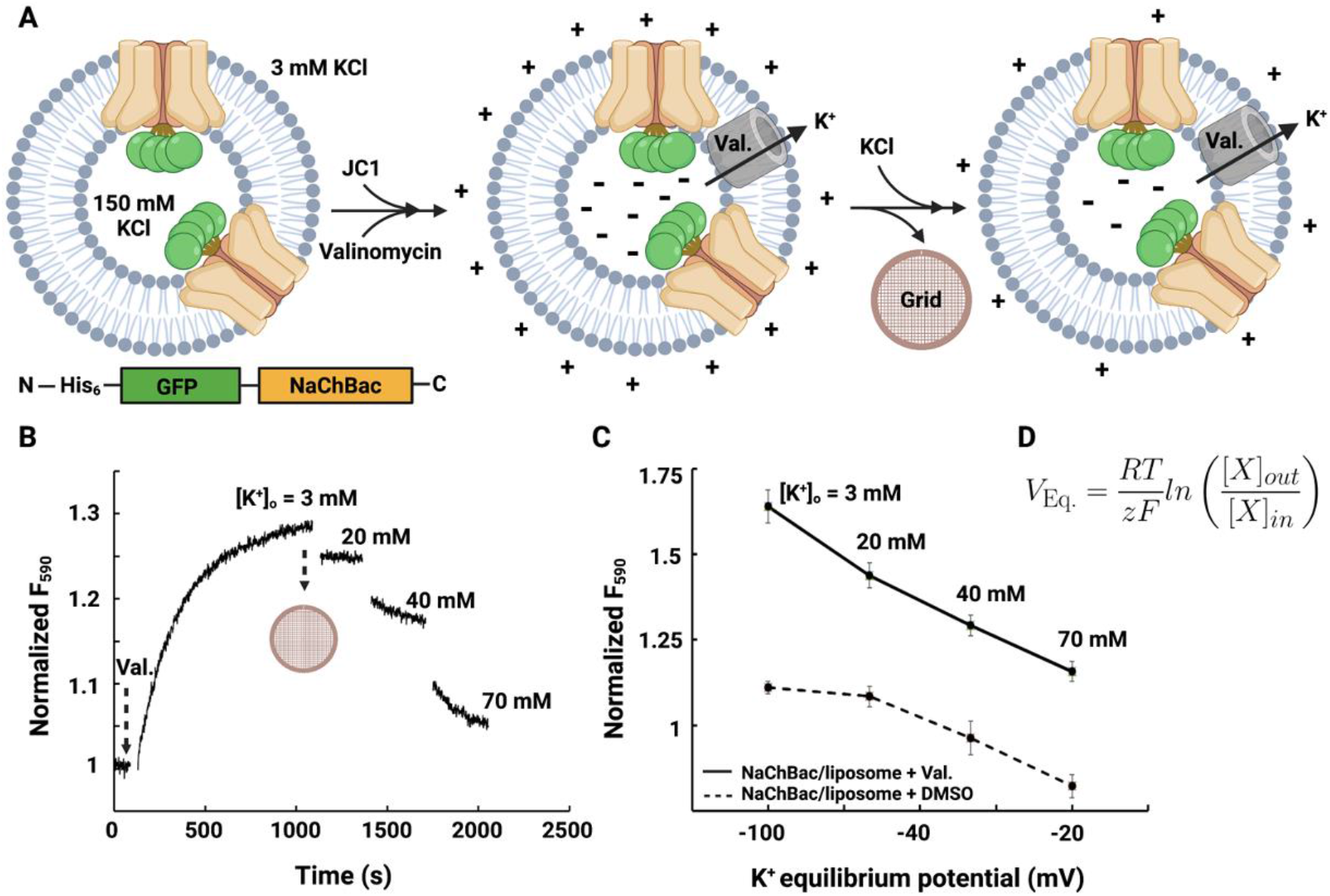
Reconstitution of NaChBac in proteoliposomes and preparation of polarized NaChBac-containing liposomes. (A) A schematic of the protocol used to obtain polarized proteoliposomes for cryo-EM analysis. Proteoliposomes loaded with 150 mM KCl were resuspended into 3 mM KCl buffer, and valinomycin was added to mediate potassium flux. Potassium efflux through valinomycin generated the negative potential inside of the vesicles with respect to the outside, and the red fluorescence of JC-1 aggregates was measured. (B) Fluorescence-based liposome flux assay of proteoliposome membrane potential. Addition of valinomycin allowed potassium efflux, and a subsequent decrease in fluorescence as external KCl concentration was increased, indicating K^+^-selective permeability and the stability of proteoliposomes. Vesicles were frozen on gold-supported gold grids for structure determination while [K^+^]_out_ = 3 mM, [K^+^]_in_ = 150 mM, membrane potential (V_Eq._) was -100 mV at room temperature (298 K). (C) Normalized fluorescence showed the difference between proteoliposomes in the presence of 1 μM valinomycin and DMSO (control), indicating potassium equilibrium potential was triggered by valinomycin. Error bars indicated s.e.m. (n = 3-5). (D) The Nernst equation. V_Eq._ is the equilibrium potential (Nernst potential) for a given ion. R is the universal gas constant. T is the temperature in Kelvin. z is the valence of the ionic species. F is Faraday’s constant. [X]_out_ and [X]_in_ are the concentration of the ionic species X in the extracellular and the intracellular fluid, respectively.

**Figure 2.**
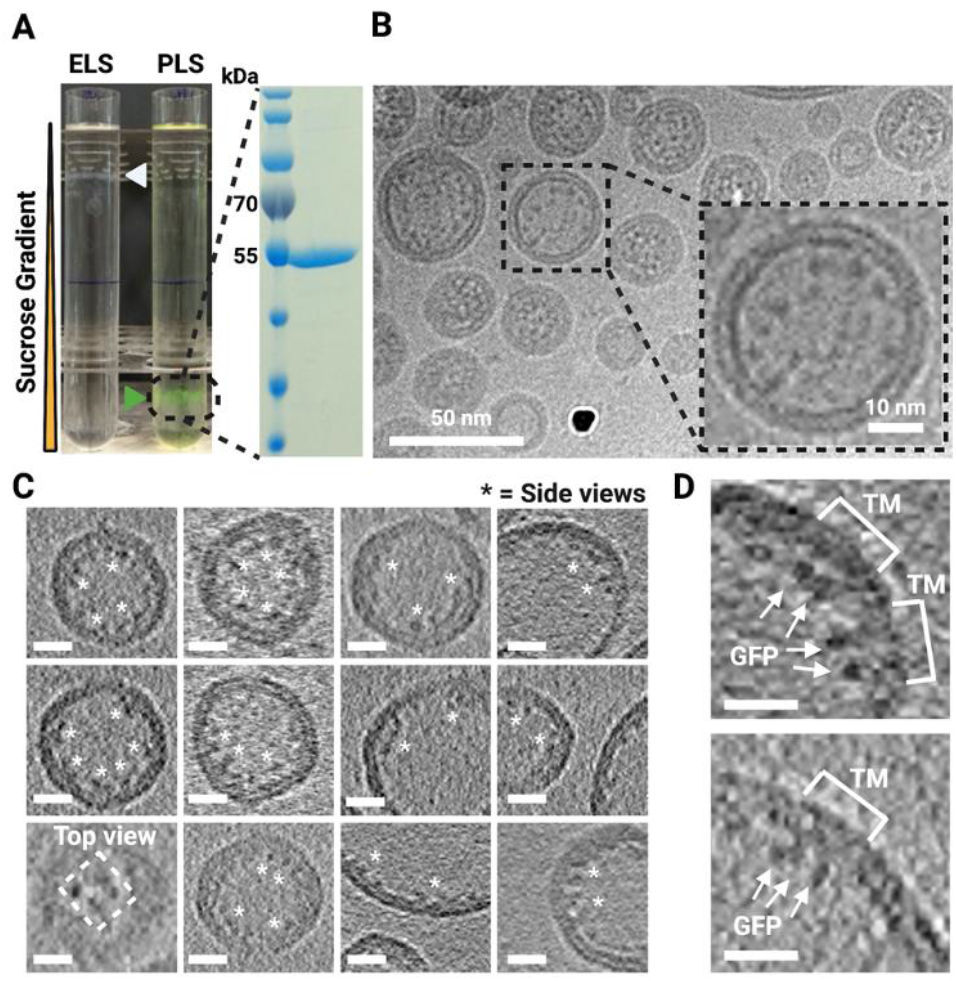
Visualizing NaChBac in proteoliposomes by cryo-EM/ET. (A) Sucrose gradient ultracentrifugation showed a significant band shift between empty liposomes (ELS) and proteoliposomes (PLS), and the protein showed no degradation on the SDS-PAGE after proteoliposome reconstitution. (B) A cryo-image of proteoliposomes. The inset showed a proteoliposome with proteinaceous densities. (C) Cryo-ET slices of proteoliposomes showed particle positions and the side view and the top view of his-GFP-NaChBac. (D) Tomographic slices of proteoliposomes showed GFPs outside the lipid bilayer and TMs of NaChBac embedded in the membrane. Scale bars: 10 nm.

### Cryo-EM/ET and Structural Analysis by StA

Proteoliposoms were screened by negative stain-EM (SI Appendix, Fig. S2) and cryo-EM showing a large population of vesicles with a diameter up to 150 nm. While some vesicles were much larger, the smaller vesicles were more amenable to producing thin ice for cryo-ET imaging. The initial cryo-EM preparations showed a significant clustering of vesicles next to the edges of the grid holes with empty hole centers. In order to have a uniform distribution of vesicles, we performed multi-sample application on gold-supported gold grids inspired by previous reports (Snijder et al, 2017; Tonggu & Wang, 2020), tuning the parameters of the sample preparation robot, resulting in the final sample for cryo-ET (Fig. 2 B).

We collected 98 tomograms (Table S1) and processed them using early versions of tomoBEAR (Balyschew et al, 2023) combining Motioncor2 (Zheng et al, 2017), Gctf (Zhang, 2016), Dynamo (Castaño-Díez et al, 2012) and IMOD (Kremer et al, 1996). Tomographic reconstructions clearly showed GFP densites and occasional densities between the membranes. The majority of the particles had the GFP “inside the vesicle”, the orientation similar to the inner membrane of bacteria. Knowledge of the orientation of the majority of the channels defined how we established the voltage across the membrane. We could observe “side-views” and tetramer-resembling “top-views” (Fig. 2 C and D). Unfortunately due to the small size of NaChBac we could not perform automated particle picking. Attempts to use geometry-supported particle identification by “drawing” membranes followed by classification of positions into protein-containing and empty membranes were unsuccessful. We therefore picked ∼86,000 tomographic positions from the cryo-ET volumes (prioritizing tomograms with thin ice) manually using the Dynamo Catalogue System.

We next performed StA. First, we used Dynamo to align all the subtomograms to a reference produced by manually aligning a small subset of particles and allowing the particles to rotate 360 degrees around the unit sphere (first two Euler angles) and applying high rotational symmetry around the vertical axis (not searching the last Euler angle). This resulted in a curved membrane with a defined density at the concave side of the membrane (Fig. 3). The membrane was less ordered at the edges of the box reflecting various sizes of the vesicles leading to “averaging out” of the bilayer. We next performed a classification of all the particles into 10 classes allowing the particles to rotate around the third Euler angle with limited rotations around the first two. This resulted in several classes resembling tetrameric features containing ∼42,100 particles, and the dataset was further cleaned to ∼31,800 particles with more classifications. Further classification based on rejecting the classes with poorly resolved membranes narrowed down the dataset to ∼25,500 particles which we took for further classification and refinement in RELION-4.0 (Zivanov et al, 2022) (SI Appendix, Fig. S3). At this point there were still significant differences between the membrane curvatures of classes. We performed several rounds of classification and tried to auto-refine the resulting datasets with more or fewer particles, and ultimately, a dataset of 3,116 particles gave us the best structure. Refinement of this dataset produced a structure at 16.3 Å resolution (Fig. 4 A and S5). During the outlined process we did not apply symmetry and only selected tetrameric-looking classes for further processing. The overview of the classification process is given in Figures 3 and S4.

**Figure 3.**
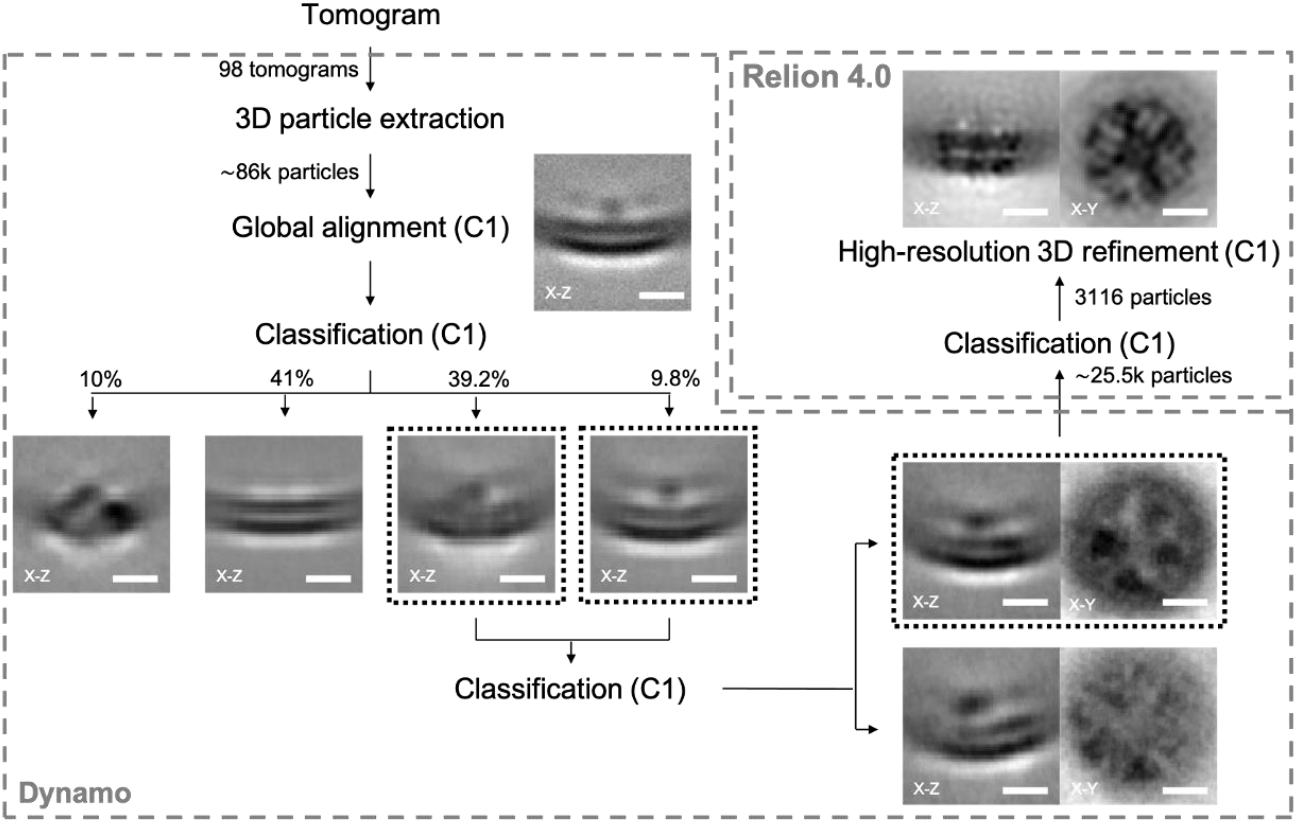
The workflow for structural determination of NaChBac embedded in liposomes by StA. Manually identified particles (∼86,000) were extracted from 98 tomograms. After a global alignment of all extracted particles, multiple rounds of classification were applied to the classes containing tetrameric features from the top-views. Particles with clear protein signals were then subjected to the RELION-4.0 for the next classification and refinement. Final subtomogram averaging from 3116 particles displayed clear domain features of NaChBac. (Scale bars: 5 nm.) More details were provided in SI Appendix, Fig. S3 and S4.

**Figure 4.**
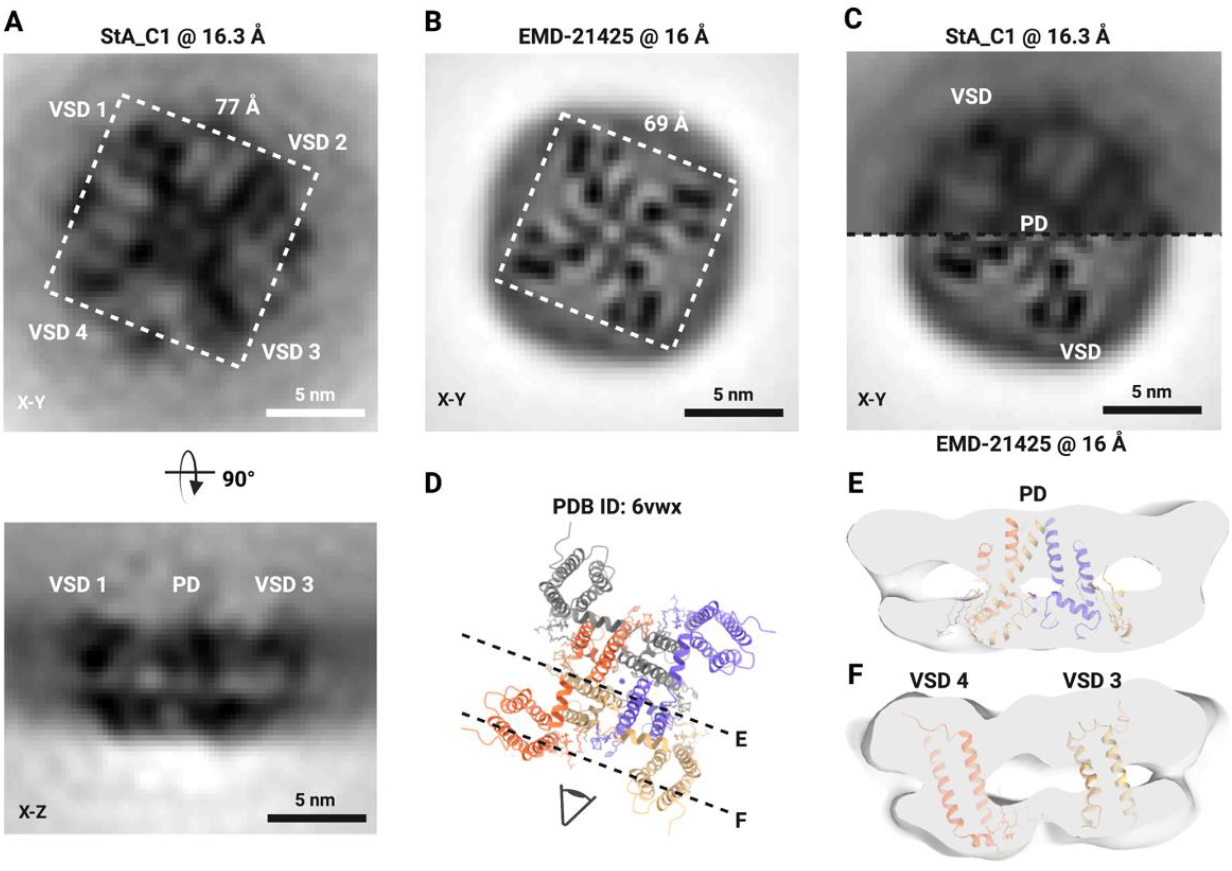
Structure of NaChBac in proteoliposomes and comparison to the structure in nanodiscs (EMD-21425). (A) StA map of NaChBac embedded in a liposome. (B) Cryo-SPA map of NaChBac embedded in nanodisc. (C) StA map (above) at 16.3 Å compared with the recalculated Cryo-SPA structure (EMD-21425) at 16 Å resolution (below). The volumes were mutually aligned. The VSD and PD could be visualized as a tetrameric feature from the top of the StA structure. (D) The atomic model of EMD-21425 in an inactivated state. (E) A segmented StA map of NaChBac embedded in a liposome overlaid with the atomic model built based on the SPA structure (EMD-21425). The model showed the density of PD surrounded by four VSDs based on the fit of the PDB ID: 6vwx to the StA map.

While the structure looked overall tetrameric, the symmetry was not perfect and refinement of the structure with C4 symmetry did not result in a better structure. Symmetry expansion also did not help to improve the structure. Comparing the structure to the intermediate class averages (Fig. 3), which we could not autorefine, we observed a heterogeneity in the positioning of the voltage-sensing domains. Finally, while we attempted multiple masks for the alignment, the best final alignment was achieved when the mask did not accommodate the densities of GFP. This could mean that the densities of GFP on a 15-long residue linker, while helpful for the localization of the proteins of interest, were flexible and did not help to align the particles to the average.

The StA map clearly showed the domains of NaChBac in the lipid bilayer. At this domain-level resolution we could identify the tetrameric-appearing pore domain (PD) which contains eight full transmembrane helices and four smaller voltage-sensing domains (VSDs), each of which contains four transmembrane helices. Comparison of the StA reconstruction with the map from the SPA structure in lipid nanodisc reported previously (EMD-21425) resampled and low-pass filtered to 16 Å showed overall similarity (Fig. 4 B and C). The corresponding atomic model fitted the StA density well, considering the moderate resolution (Fig. 4 D and E). Interestingly, the StA structure was ∼8 Å, or ∼10% wider compared to the structure in nanodiscs filtered to the same resolution (Fig. 4 A and B).

## DISCUSSION

We showed that in principle it is possible to identify and align small and mostly transmembrane proteins using StA and to obtain a correct structure, although at a modest resolution. Here, we could use the fused GFP in order to locate the proteins of interest in tomograms and to perform a rough alignment, but for the final steps of the alignment the GFP was not useful, due to the flexibility of the linker region. Perhaps a more ordered interacting partner such as a specific toxin (Xu *et al*, 2019), a Fab fragment (Wu et al, 2012) or a legobody (Wu & Rapoport, 2021) would be more useful to support particle alignment to the average. We did not use toxins or Fab fragments as those structures would be likely similar to the previously reported structures of purified channels (Xu *et al*, 2019; Wisedchaisri *et al*, 2021). The tetrameric NaChBac with a molecular weight of ∼120 kDa with mostly transmembrane residues is probably on the smaller side of the molecular weight that could be approached using the current technology. For our structural analysis we imaged the channels in thin ice, picked a large number of particles and used a combination of classification algorithms (Dynamo and RELION-4.0). Even with such optimizations, we could only obtain the resolution of ∼16 Å. While it is higher than the average resolution of StA structures deposited to the Electron Microscopy Data Bank in 2022 (20 Å), it only provides domain-level resolution. We believe that imaging of even smaller proteins in membranes without large soluble domains such as monomeric GPCRs without interaction partners may be too ambitious at the current level of technology.

At the obtained resolution we could reliably observe the pore and the voltage-sensing domains of the channel. To this end we could estimate that the structure of the channel in proteoliposomes is ∼10% wider than the structure reported in nanodiscs (EMD-21425) filtered to the same resolution. This could be physiologically relevant for two reasons. Firstly, nanodiscs were shown to confine the lipids inside the polymer (Schachter et al, 2020) and to modify the conformations of pentameric ligand-gated ion channels (Dalal et al, 2022). Secondly, molecular dynamics simulations for the K_v_ channel showed substantial motion of the VSDs (Jensen *et al*, 2012) that on average could make the proteins wider compared to the compact versions that are typically observed in nanodiscs. The possible motion of the VSD domains is another factor that potentially limits the obtainable resolution and could partially explain that the application of the C4 symmetry in our case did not lead to a much improved structure. While the observed increased protein diameter might be a result of the applied membrane potential, a recent report by Mandala and MacKinnon on the movements of the VSDs the Eag K_v_ channel showed that the cryo-EM structures of the channel with and without the transmembrane potential were overall similar in size. Therefore it is possible that in the case of NaChBac, the increased size of the channel is not a consequence of the applied potential but as a result of imaging of the channel in a lipid bilayer without physical restraints.

With future improvements in detectors (Ruskin et al, 2013; Wu et al, 2016; Nakane et al, 2020) and phase plates (Schwartz et al, 2019) they may be possible to get higher quality data and consequently higher resolution of NaChBac or proteins of similar sizes in proteoliposomes. Higher number of particles could improve the resolution; they may be obtained by processing more tomograms and picking more protein copies for StA. While the reconstruction of tomograms can be streamlined (Mastronarde & Held, 2017; Balyschew et al, 2023), we did not manage to automate particle picking and had to pick them manually which is a very laborious process. Developments in machine learning and the potential applications for particle picking (de Teresa-Trueba *et al*, 2023; Rice *et al*, 2022; Zeng *et al*, 2023) could streamline particle picking in the future. Our dataset will be on the difficult side for such applications.

## MATERIALS AND METHODS

### Expression and Purification of NaChBac

The cDNA for full-length NaChBac was cloned into pET21a with an amino terminal His_6_-GFP^A206K^. Overexpression in *E. coli* C41(DE3) cells was induced with 0.25 mM IPTG (final concentration) at 22 °C when the OD_600_ reached 0.8 to 1.0. Cells were harvested after 20h incubation at 22 °C, and cell pellets were resuspended in a buffer containing 25 mM Tris, pH 8.5, and 300 mM NaCl. Cells were disrupted by high pressure homogenizer, and insoluble fractions were removed by centrifugation at 30,000 × g for 30 minutes. The supernatant was subjected to ultracentrifugation at 150,000 × g for 2 hours. The membrane-containing pellets were resuspended in extraction buffer containing 25 mM Tris, pH 8.5, 300 mM NaCl, 20 mM imidazole, and 1% (wt/vol) DDM, incubated at 4 °C for 2 hours, and subsequently centrifuged at 30,000 × g for 30 minutes. The supernatant was applied to Ni-NTA resin, washed with 20 column volumes of wash buffer containing 25 mM Tris, pH 8.5, 300 mM NaCl, 20-80 mM imidazole, and 0.1% DDM. Target proteins were eluted with 3 column volumes of elution buffer containing 25 mM Tris, pH 8.5, 300 mM NaCl, 250 mM imidazole, and 0.1% DDM. After concentration, proteins were further purified by SEC (Superose 6 Increase 10/300 GL; GE Healthcare) in a running buffer containing 25 mM Tris, pH 8.5, 150 mM KCl, and 0.1% DDM.

### NaChBac Reconstitution into Liposomes

The protocol for proteoliposome reconstitution was carried out as described before (Geertsma et al, 2008), with minor modifications. *E. coli* Polar Lipid Extract (Avanti) dissolved in 400 μL chloroform at 25 mg/mL was dried to a thin film under a gentle stream of nitrogen and resuspended in 1 mL reconstitution buffer containing 25 mM HEPES, pH 7.0, and 150 mM KCl. After water-bath sonication for 5 minutes, the lipid solution was frozen with liquid nitrogen and thawed in water bath for 10 times, and the lipid solution was subject to repeated extrusion through 100 nm filters. Destabilization of liposomes was incubated with 2% (wt/vol) DDM (Anatrace) as a final concentration at 25 °C for 2 hours. Then purified NaChBac was added to make the lipid-to-protein ratio of 2:1 (wt/wt). After incubation at 4 °C for 1 hour, the lipid-protein-detergent mixture was loaded into dialysis bag against the reconstitution buffer at 4 °C for approximately one week with gentle rotation, and Bio-Beads SM-2 resin (Bio-Rad) 0.4 g was then added to remove residual detergents from the proteoliposomes. After incubation at 4 °C overnight, Bio-beads were removed through filtration, and the proteoliposomes were ready for liposome flux assays.

### Density Gradient Centrifugation

Sucrose gradients were prepared in SW 40 Ti ultracentrifuge tubes (Beckman Coulter) on a Biocomp Gradient Master (ScienceServices, Mnchen) based on the method of Coombs and Watts (Coombs & Watts, 1985). Concentrated sucrose solution (25 mM HEPES, pH 7.0, 150 mM KCl, and 1 M sucrose) was layered under an equal volume of light solution (25 mM HEPES, pH 7.0, 150 mM KCl, and 0.3 M sucrose) in centrifuge tubes. The tubes were closed with caps to expel all air, and the gradient (10-30%) was formed by rotation. A 400 mL volume was removed from the top of each tube before the sample was added, and a 400 mL liposome with or without protein per tube was loaded and centrifuged at 130,000 x g for 18 hours at 4 °C. After ultracentrifugation, opaque liposome bands were collected with a syringe and diluted with reconstitution buffer to remove most of the sucrose. Liposomes were pelleted at 90,000 x g for 1 hour at 4 °C and resuspended in the reconstitution buffer, and samples were identified by SDS-PAGE.

### Negative-staining EM

Proteoliposome samples were diluted in a buffer containing 25 mM HEPES, pH 7.0, and 150 mM KCl to a series concentration. The sample was adsorbed to freshly glow-discharged carbon-coated grids, rinsed with several drops of the dilution buffer, and stained with 1% uranyl acetate. Images were recorded at a magnification of 49,000 × with defocus values ranging from -2.5 to -3.5 µm, resulting in a pixel size of 2.26 Å/pixel (Spirit Biotwin, FEI). As most of the protein was inserted with GFP inside it was not possible to screen protein insertion with negative staining EM and had to be done with cryo-EM.

### Liposome Flux Assay

The proteoliposome vesicles prepared in 150 mM KCl were diluted 100-fold into 3 mM KCl solution (25 mM HEPES, pH 7.0, and 3 mM KCl) containing 1 μM of JC-1 (5P,5P,6,6P-tetrachloro-1,1,3,3P-tetraethylbenzimadazolylcarbocyanine iodide, Invitrogen). After the JC-1 fluorescence stabilized, valinomycin was added into the system to final concentration of 1 μM. The fluorescence signal of the J-aggregates (λ_ex_ = 480 nm, λ_em_ = 595 nm) was monitored when valinomycin solution initiated K^+^ efflux from all the vesicles until the external K^+^ concentration was increased by the addition of 2.5 M KCl solution. The normalized data were averaged across three independent measurements and the mean and SDs were reported.

### Preparation of Polarized Lipid Vesicles and Cryo-EM Grids

The proteoliposomes prepared above were diluted into 10-nm gold fiducial markers in 3 mM KCl solution (25 mM HEPES, pH 7.0, and 3 mM KCl), and 1 μM valinomycin was added and incubated for 5 minutes on ice. An aliquot (2 μL) of this polarized vesicle solution was applied onto a glow-discharged holey gold grid (Quantifoil Au R2/2, 400 mesh). After incubating the sample on the grid for 3 minutes at 10 °C with a humidity of 100%, the grid was manually blotted from the side using a filter paper. Another 2 μL of the polarized vesicle solution was applied to the same grid for 15 seconds, and then the grid was blotted with Whatman^®^ No. 1 filter paper and plunge-frozen in liquid ethane (Vitrobot Mark IV, Thermo Fisher Scientific).

### Cryo-ET Data Collection, Image Processing, and Subtomogram Averaging

Imaging was performed on a Titan Krios G2 Cryo-TEM (Thermo Fisher Scientific) with a K3 direct detection camera (Gatan) and a BioQuantum imaging filter slit width of 20 eV (Gatan) operated by SerialEM software (Schorb et al, 2019). Tomographic series were acquired using a dose-symmetric scheme (Hagen et al, 2017), with tilt range ±45°, 3° angular increment and defoci between -2.5 and -3.5 μm. The acquisition magnification was 81,000 ×, resulting in a calibrated pixel size of 1.39 Å. The electron dose for every untilted image was increased to around 20 e^-^/Å^2^, and tilt images were recorded as 10-frame movies in counting mode and a total dose per tilt series of around 130 e^-^/Å^2^.

Data processing was performed using early versions of tomoBEAR (Balyschew et al, 2023) implemented as a set of Matlab scripts. Frames were aligned and motion-corrected using MotionCor2 (Zheng et al, 2017). Tilt series were aligned using 10-nm gold fiducial markers by IMOD (Kremer et al, 1996; Mastronarde & Held, 2017). Contrast transfer function (CTF) estimation was performed using defocus values measured by Gctf (Zhang, 2016) for each projection. A total of 98 tomograms and the four binned reconstructions were generated from CTF-corrected, aligned stacks using weighted back projection in IMOD. Subtomogram positions (∼86,000) were picked manually from 4-times binned tomograms and extracted with a box size of 128 cubic voxels from binned 2 tomograms using the Dynamo Catalogue system (Castaño-Díez *et al*, 2017). Initial alignment was done manually on ∼200 particles, and the center of four GFPs and the direction of the central axis based on the membrane were defined using *dynamo_gallery* after which the roughly aligned particles were summed up low-pass-filtered to 40 Å. This volume was used as an initial reference for the global alignment of all subtomograms, resulting in all particles to the same Z height in 3D. Several rounds of initial classification by multireference alignment were used to remove junk particles, with the first step 360° in-plane search (XY plane) being performed on all particles in C1, and a large soft-edged sphere mask was applied for the particle cleaning. Further multireference alignment and averaging with a soft-edged ellipsoid alignment mask was applied throughout and the averaging results showed a prominent tetrameric feature in between the lipid bilayer in C1. In total, ∼25,500 particles from good class averages were then subjected to the RELION-4.0 (Zivanov et al, 2022) for further 3D classification and refinement. Final converged averages were formed by 3116 particles in C1 at 16.3 Å resolution, and a smaller sphere mask was applied to improve the resolution. No symmetry was applied during processing.

## Acknowledgments

We thank the Kudryashev Group members Nikita Balyschew, Kendra E. Leigh, Ricardo M. Sanchez, and Vasilii Mikrtumov for useful discussions and advice on data processing. We thank Eric Geehrstma, Klaus Fendler and Werner Kühlbrandt for useful discussions. The work was funded by Scholarship from the G-max Precision Co., Ltd. and Taiwanese Government Scholarship to Study Abroad to Shih-Ying Scott Chang, a DFG Project grant to KU 3221/2-1, the Heisenberg Award KU3221/3-1, the Sofja Kovalevskaja Award from the Alexander von Humboldt Foundation to Misha Kudryashev. We thank Juan Castillo-Hernandez and Oezkan Yildiz from the Max Planck Institute for Biophysics for IT support. We thank Deryck Mills, Sonja Welsch and the EM staff from the Max Planck Institute for Biophysics for expert support during cryo-ET data collection.

## Author contributions

S.-Y.S.C. performed protein expression, purification, established proteoliposome reconstitution and the membrane potential assays, performed cryo-grid optimization, cryo-ET imaging, and data analysis; wrote the manuscript with M.K. S.A.W. performed initial protein expression and purifications, designed and supervised by P.M.D. M.K. designed and supervised the project, obtained funding, and wrote the manuscript with S.-Y. S. C.

## Data and materials availability

The structure and the original data will be deposited to EMDB and EMPIAR and the accession codes will be stated here.

