## Supplementary Material for "Determining The Structure of the Bacterial Voltage-gated Sodium Channel NaChBac Embedded in Liposomes by Cryo Electron Tomography and Subtomogram Averaging"

### SI Appendix

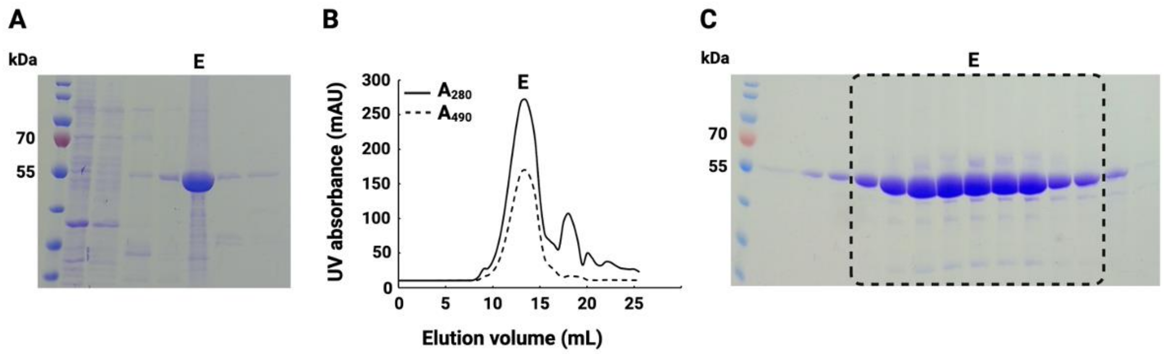


***Figure S1 Purification of NaChBac.*** *(A) SDS-PAGE of protein purification with Ni-NTA column. E, elution. (B and C) SEC purification of NaChBac in DDM detergent with a monodispersed peak. E, elution.*

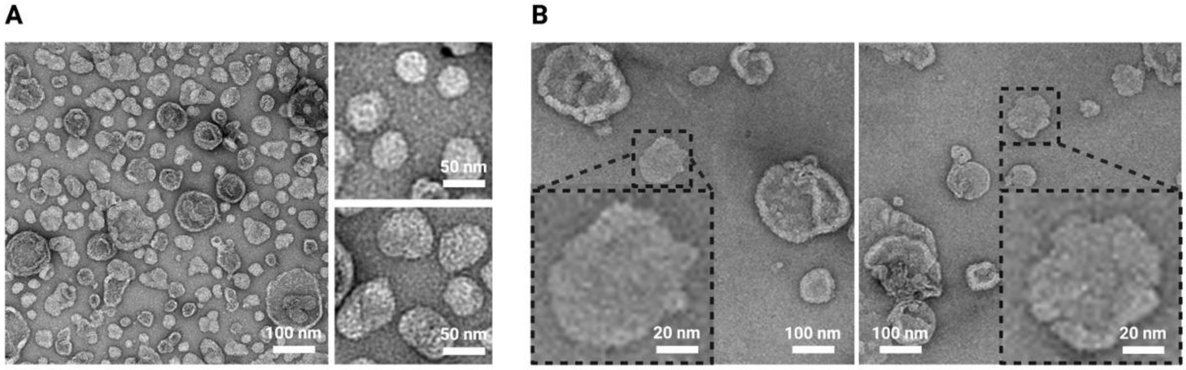


***Figure S2 Negative-staining images of proteoliposomes.*** *(A) A negative-staining image of proteoliposomes with detergent removal entirely. The insets showed proteoliposomes with "sharp edge" features. (B) Negative-staining images of proteoliposomes with residual detergent during dialysis. The insets showed proteoliposomes with "fluid edge" features.*

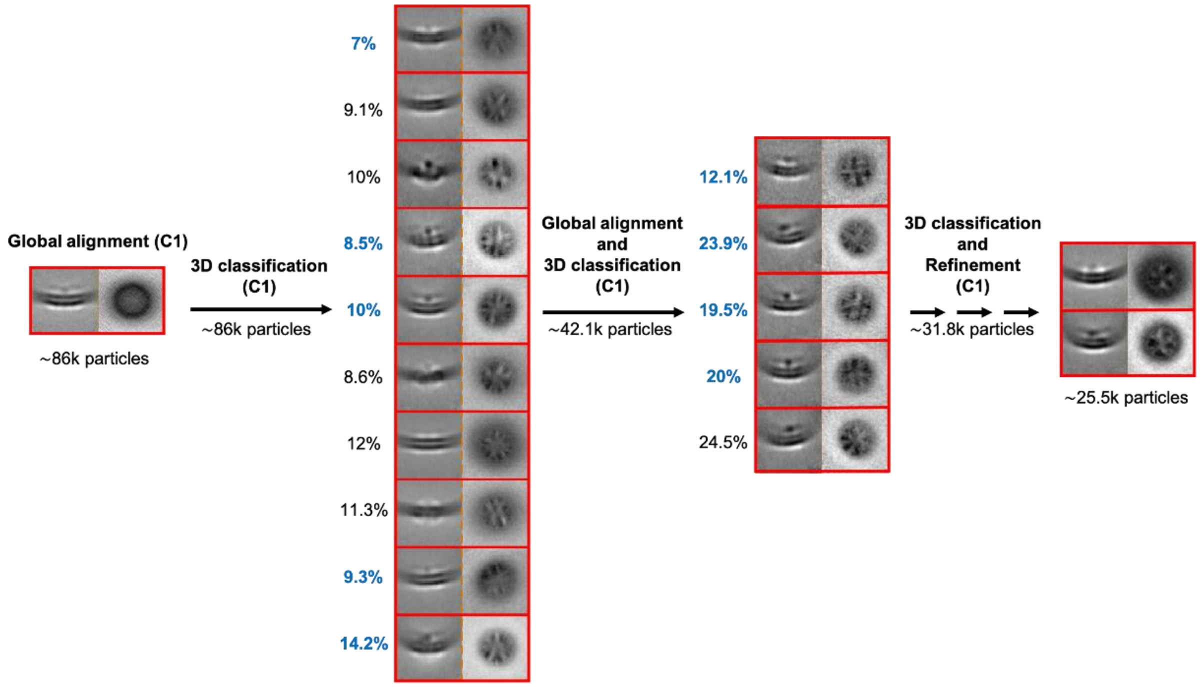


***Figure S3 Strategy of classification for the dataset cleaning with Dynamo in C1.*** *˜86,000 particles were aligned globally and were implemented to 3D classification for the particle cleaning. ˜42,100 particles were selected to perform further classification. After collecting good particles (˜31,800) from previous steps, multiple rounds of 3D classification and refinement were performed, and ˜25,500 particles were then subjected to the RELION-4.0 for further 3D classification and refinement.*

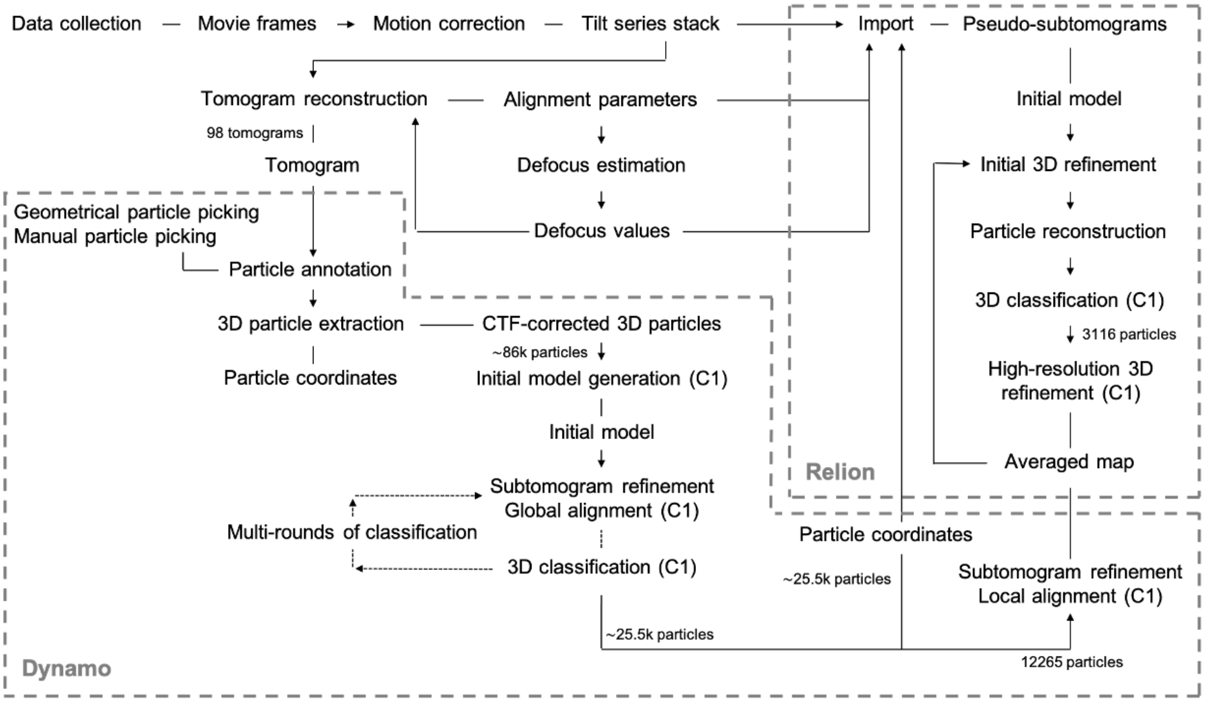


***Figure S4 Workflow for tomogram reconstruction and subtomogram averaging.*** *Tilt series stack alignment, tomographic reconstruction, particle annotation, 3D classification, and subtomogram averaging were shown in the workflow.*

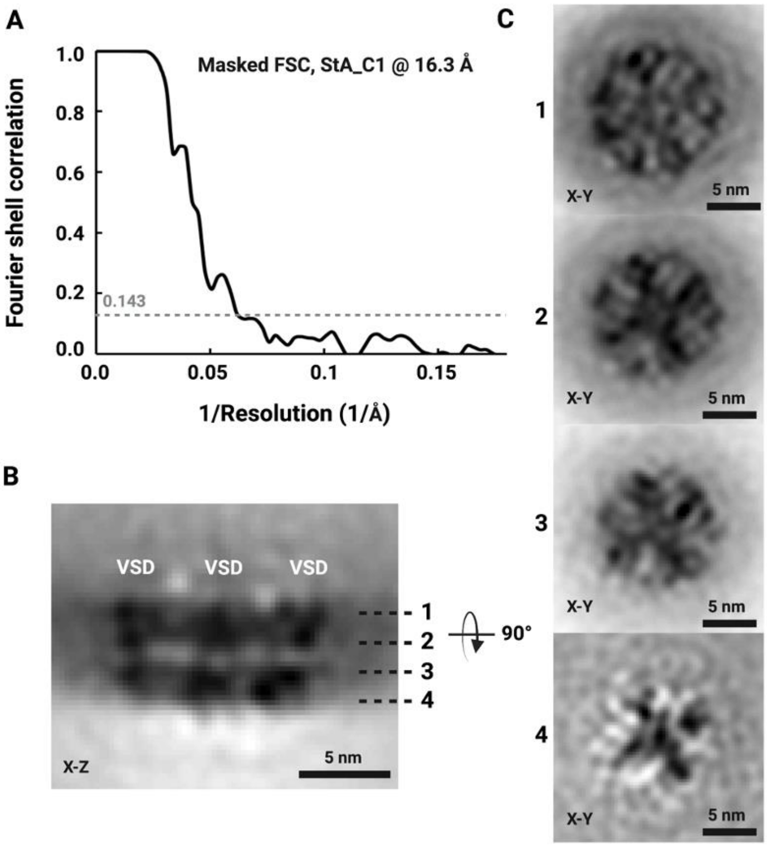


***Figure S5 FSC curve of the masked map and slices through the structure of NaChBac map in liposomes along the z-axis.*** *(A) Gold standard FSC (criteria 0.143) curve of the masked map with a reported resolution of 16.3 Å for the map with C1 symmetry. (B and C) The VSD and PD could be visualized as a tetrameric feature in the X-Y plane along the z-axis.*

**Table S1 Data Processing Statistics**

| | **NaChBac in liposomes; EMD-17163** |
|---|---|
| **Data collection and processing** | |
| Microscope | Titan Krios G2 |
| Magnification | 81,000 x |
| Voltage (kV), Cs | 300 kV, 2.7 mm |
| Total electron dose ($e^-/A^2$) | ˜130 |
| Defocus range (µm) | -2.5 to -3.5 |
| Camera | Gatan K3 + BioQuantum image filter |
| Pixel size (Å) | 1.393 |
| Number of tomograms | 98 |
| Symmetry imposed | C1 |
| Initial particles | ˜86,000 |
| Final particles | 3,116 |
| Refinement method | Independent half-set |
| Map resolution (Å) | 16.3 |
| FSC threshold | 0.143 |
